# Meta-analysis of Scandinavian Schizophrenia Exomes

**DOI:** 10.1101/836957

**Authors:** Francesco Lescai, Jakob Grove, F. Kyle Satterstrom, Elliott Rees, Jonas Bybjerg-Grauholm, Thomas Damm Als, Jaroslaw Kalinowski, Anders Halager, Tarjinder Singh, Qibin Li, Jun Wang, James T R Walters, Michael J Owen, Michael C O’Donovan, Merete Nordentoft, Preben Bo Mortensen, David M Hougaard, Thomas Werge, Ole Mors, Benjamin M Neale, Mark J Daly, Anders D Børglum

**Author notes:** Correspondence: Francesco Lescai and Anders D Børglum.

## Abstract

Rare genetic variants may play a prominent role in schizophrenia. We report on the to date largest whole exome sequencing study of schizophrenia case-control samples from related populations and combine with other available sequence data, analysing in total 34,084 individuals (14,302 cases). Three genes showed significant association at FDR < 0.10 (*SETD1A*, *TAF13* and *MKI67*) and gene-set analyses highlighted the involvement of the synaptome and excitatory neurons, and demonstrated shared architecture with high-functioning autism.

## Main text

Schizophrenia is one of the most disabling psychiatric disorders. With heritability estimates of 60-80%^1,2^ genetics is the most prominent etiologic component and the genetic architecture is extremely polygenic including both common and rare variants^3–5^. Common variants have been estimated to explain ¼-1/3 of the heritability^3,6^, leaving a large proportion of the liability unaccounted for and indicating that rare variants may play a substantial role as recently observed in other complex traits^7^.

Schizophrenia has a markedly reduced fecundity suggesting that variants with large effect sizes will swiftly be removed from the population by purifying selective pressure^8^. Accordingly, most rare variants identified so far are large copy number variants occurring de novo in affected offspring^9^ and only a single gene, *SETD1A*, harbouring deleterious de novo point mutations, has been robustly identif^10,11^. Whole exome sequencing studies of rare variants in case-control samples have found an increased burden of particularly ultrarare protein-disrupting or -damaging variants (dURVs) among cases, identifying gene sets enriched for these variants and demonstrating that larger sample sizes are needed to identify the specific risk genes hit by dURVs^4,5,12^.

Here we investigate the largest case-control sample of schizophrenia exomes to date, from a confined geographic region in Scandinavia with genetically^13^. To empower gene discovery, we combine the case-control results with results on published de novo mutations, and follow-up the top-findings in case-control exome data from the UK10KConsortium^14^ as well as targeted sequence data^15^. In total we analyse sequence data from 34,084 individuals, including 14,302 cases.

First, we set to analyse with the same methods three datasets from Scandinavian populations: a previously investigated Swedish sample (dbGAP phs000473.v2.p2, 4,969 cases and 6,245 controls^5^), a Danish cohort from iPSYCH^16^ (3,710 cases and 6,375 controls) and a second Danish cohort recruited in clinical centres (1,866 cases and 1,093 controls). All samples were filtered to exclude relatedness, population structure, and outliers in the distribution of ultra-rare variants (Online Methods 1.2, 1.3 and 2.1). The analyses were performed on each dataset, and a random-effect meta-analysis was conducted to combine the results, in a total sample of 7,744 cases and 11,176 controls (Supplementary Figure S1). Based on results from previous exome studies and similar results from the present exomes (Online Methods 2.4), we focused on ultra-rare variants (URVs), defined as singletons in the dataset under analysis and absent from the ExAC database, after excluding samples with psychiatric disorders. The variants were further classified as synonymous, missense non-damaging, damaging and disruptive according to previous studies (Genovese et al. ^5^, and Online Methods 2.5, Supplementary Figure S2). First, we investigated the burden of different URV categories on loss-of-function intolerant genes^17^: the analysis was conducted by doing logistic regression on the count of URV for each individual and using a large number of covariates to account for potential confounding factors (Online Methods 3.1 and 3.2); the results (Figure 1) showed a significant burden of disruptive URVs (1.54*10^−13^ p-value for the meta-analysis). A similar trend was observed for missense damaging URVs, although the burden was significant only in the Swedish dataset, supporting that damaging and disruptive URVs (dURVs) can be combined together, as previously reported^5^. The burden of dURVs in Loss of Function (LoF) intolerant genes showed a p-value of 4.89*10^−6^. A similar association was observed for disruptive, damaging and dURVs also in missense-constrained genes (Supplementary Figure S3). We selected dURVs as our primary analysis. We followed up the genome-wide burden analysis, by regressing on the counts of dURVs for a number of gene-sets previously identified based on pathways and expression analysis^5^: the results of this analysis (Supplementary Figure S4) showed a significant increase in genes annotated in the synaptome (p-value 8.3*10^−3^), and in genes regulated by CHD8 (1.5*10^−4^), a transcription factor previously implicated in autism^18,19^. Interestingly, among 102 autism spectrum disorder (ASD) risk genes recently identified by the Autism Sequencing^20^, the 53 *ASD predominant* genes showed a significant genome-wide burden, driven by disruptive URVs (p-value 9.07*10^−4^) (Supplementary Figure S4 and Supplementary Table 1), while the 49 genes enriched for variants in ASD cases with severe neurodevelopmental delay (*ASD NDD* genes) did not show any increased burden. The regression analysis with the counts of synonymous URVs on the same sets, performed as control, showed no significant results (Figure S4).

**Figure 1.**
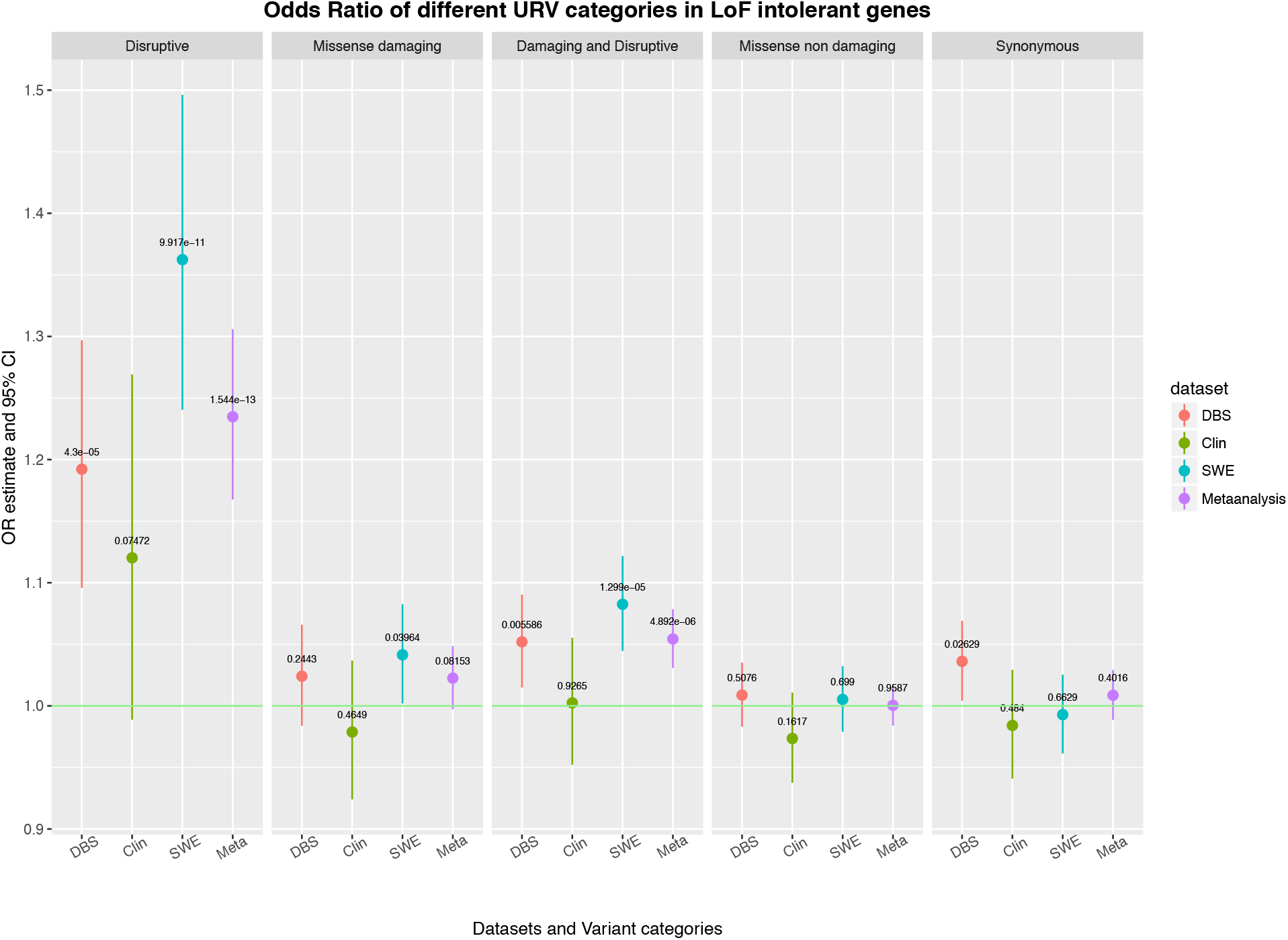
Odds Ratio estimates and 95% confidence intervals for genome-wide enrichment of different variant annotations in loss-of-function constrained (pLI>0.9) genes in each of the dataset, and random-effects meta-analysis. Datasets: Danish Biobank dried blood spots (DBS), SCZ clinical centres in Denmark (Clin), Swedish dataset from Ganna et al. (SWE), random-effects meta-analysis (metaanalysis).

The analysis of individual genes was performed with SKAT (Online Methods 5 and 6, and Supplementary Figures S5 and S6), and results of dURV analysis are shown in Supplementary Table 2 with the top ranking genes *EFCAB4B* (CRACR2A P = 2.4*10^−4^) and *DLGAP4* (P = 3.2*10^−4^) encoding, respectively, calcium release activated channel regulator 2A and DAP-4 which is a signaling molecule found at the postsynaptic density in neuronal cells that can interact with potassium channels and receptors. Although not surpassing genome-wide significance it is of note that both genes have been linked with schizophrenia or other mental disorders^21,22^.

To increase the power for gene discovery we jointly analyzed our case-control dataset together with de novo mutation data from 1,077 case-parents trios^23^, using an extended version of the Transmission And De novo Association (extTADA) test developed by H. Nguyen and colleagues^24^. As recommended by the authors, we clustered the data in different subsets, based on their covariates matrix in order to minimise the effect of covariates, which cannot be accounted for in the model itself (Online Methods 7.1). We identified subset of data with high correlation between the results of the genome-wide URV burden analysis with and without covariates (Supplementary Figure S7) and used these subsets as datasets to be combined in the extTADA analysis. The results suggest that 5.4% of genes in our sample contribute to the risk of schizophrenia (i.e. approximately 1,000 genes). In this analysis, we also found that the mean relative risk for dURVs is 1.23 (Supplementary Figure S8). Three genes showed significant association at FDR 0.10, including *SETD1A*, *TAF13* and *MKI67* (Table 1). These results lend further support to *SETD1A* and, by adding evidence to previous reports^5,23,24^, suggest *TAF13* and *MKI67* as genes conveying risk of schizophrenia if hit by dURVs. *TAF13*, encoding a transcription initiation factor, is involved with neuronal development and homozygous *TAF13* missense mutations may cause a Mendelian recessive form of intellectual disability and microcephaly^25^. *MKI67* may, in addition to functions in regulating the cell cycle, have specialized functions in the adult nervous system^26^.

**Table 1:** Genes at q-value < 0.2, in the analysis performed with extTADA. The results are annotated with the gene function, as reported in WikiGene

| Gene | BF | PP | q value | Gene Description |
| --- | --- | --- | --- | --- |
| SETD1A | 3525.40 | 0.99 | 0.00498 | SET domain<br>containing 1A |
| TAF13 | 293.20 | 0.94 | 0.03088 | TATA-box binding<br>protein associated<br>factor 13 |
| MKI67 | 81.60 | 0.82 | 0.07987 | marker of<br>proliferation Ki-67 |
| EPHA2 | 64.64 | 0.79 | 0.11352 | EPH receptor A2 |
| ZDHHC5 | 41.89 | 0.70 | 0.15010 | zinc finger DHHC-type containing 5 |
| CEP104 | 27.50 | 0.61 | 0.19024 | centrosomal protein<br>104 |

Among the six top genes (FDR<0.20, Table 1), *SETD1A* and *MKI67* harbour dURVs in the UK10K exome dataset (1,352 cases and 4,769 controls)^14^, and for both genes the same direction of effect was observed (Supplementary Table 3). In a targeted sequencing study (5,207 cases and 4,991 controls^15^), the single dURV seen was in *MKI67*, among the controls. The scarcity of dURVs in the top-ranking genes in the UK cohorts indicates that studies of ultrarare variants in closely related populations may be an advantageous approach. Combining the results on *SETD1A* and *MKI67* from all datasets with a weighted Z-method yielded a p-value of 1.36*10^−7^ and 1.87*10^−4^ for the two genes, respectively (Online Methods 8.2).

Finally, a gene-set enrichment analysis performed with hypergeometric Fisher test on the top-100 extTADA associations confirmed the results of the genome-wide regression analysis, by highlighting the importance of CHD8 regulated genes, synaptome genes, as well as genes expressed in excitatory neurons (Figure 2, Online Methods 9 and full results in Supplementary Table 4`). The results also reinforced the enrichment of ASD-predominant genes while the ASD-NDD genes showed no enrichment, suggesting that the (rare variant) genetic architecture of schizophrenia is primarily shared with high-functioning ASD and not (or to a lesser extent) with ASD ascertained for neurodevelopmental delay.

**Figure 2.**
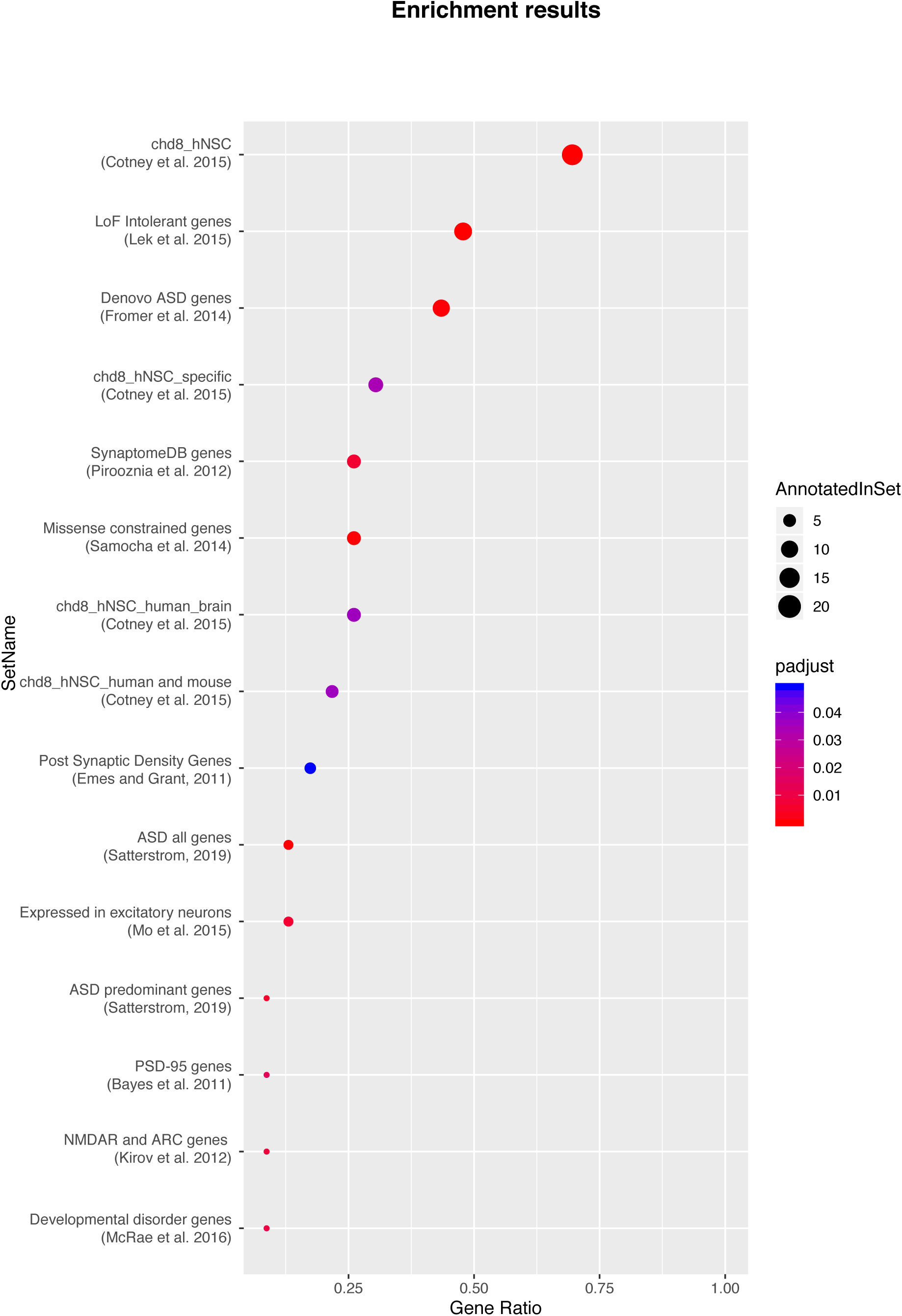
Enrichment results of top-100 significant genes in the extTADA analysis. Gene Ratio represents the ratio of significant genes belonging to the set compared to total significant genes. “padjust” indicates the FDR adjusted p-value (Benjamini-Hochberg method).

The results presented underline the role of rare variants in schizophrenia, reveal novel aspects of the genetic architecture, and suggest that larger studies are needed to identify more of the specific risk genes and involved biology.

## Acknowledgements

The iPSYCH project is funded by the Lundbeck Foundation (grant numbers R102-A9118 and R155-2014-1724) and the universities and university hospitals of Aarhus and Copenhagen. The Danish Neonatal Screening Biobank resource at the Statens Serum Institut was supported by the Novo Nordisk Foundation. Sequencing of iPSYCH samples was supported by grants from the Simons Foundation (SFARI 311789 to M.J.D.) and the Stanley Foundation and from the Lundbeck Foundation (Sequencing Schizophrenia to A.D.B.). Other support for this study was received from the NIMH (5U01MH094432-02, 5U01MH111660-02, and U01MH100229 to M.J.D.). Computational resources for handling and statistical analysis of iPSYCH data on the GenomeDK and Computerome HPC facilities were provided by, respectively, Centre for Integrative Sequencing, iSEQ, Aarhus University, Denmark (grant to A.D.B.), and iPSYCH. The Swedish datasets used for the analysis described in this manuscript were obtained from dbGaP at http://www.ncbi.nlm.nih.gov/gap through dbGaP accession number phs000473.v2.p2. Samples used for data analysis were provided by the Swedish Cohort Collection supported by the NIMH grant R01MH077139, the Sylvan C. Herman Foundation, the Stanley Medical Research Institute and The Swedish Research Council (grants 2009-4959 and 2011-4659). Support for the exome sequencing was provided by the NIMH Grand Opportunity grant RCMH089905, the Sylvan C. Herman Foundation, a grant from the Stanley Medical Research Institute and multiple gifts to the Stanley Center for Psychiatric Research at the Broad Institute of MIT and Harvard.

## Online Methods - Meta-analysis of Scandinavian Schizophrenia Exomes

### 1. The study sample

#### 1.1 iPSYCH Sample

This sample (denoted “DBS” in figures and tables) is nested in the iPSYCH2012 sample, which is a population based case-cohort study sampled from a baseline cohort consisting of all children born in Denmark between May 1st 1981 and December 31st 2005^1^. Cases were identified in the Danish Psychiatric Central Research Register^2^, that holds data on all individuals treated in Denmark at psychiatric hospitals and from outpatient psychiatric clinics. Cases in iPSYCH2012 were identified with diagnoses of schizophrenia, bipolar affective disorder, affective disorder, ASD and ADHD up until 2012. The control cohort constitute a random sample from the set of eligible subjects. DNA was extracted from Guthrie cards stored at the Danish Newborn Screening Biobank at Statens Serum Institute. For exome sequencing, we selected a subset of about 20,000 ancestry-matched samples that were sequenced by the Genomics Platform of the Broad Institute in Cambridge, MA, using an Illumina Nextera capture kit and an Illumina HiSeq.

For this study we selected 3,710 cases with schizophrenia and 6,375 controls from the control cohort with no diagnosis of schizophrenia.

#### 1.2 Clinical Sample

This sample (denoted “CLIN” in figures and tables) included 1,866 and 1,093 controls. The patients were recruited from psychiatric hospitals in the Central Denmark and the Capital regions and diagnosed with schizophrenia according to ICD 10. The control individuals were ethnically Danish blood donors.

The library preparation was performed according to the manufacturer’s instructions, and the exome was captured using Agilent SureSelect version 3 (Agilent Technologies, Santa Clara, CA, USA). The libraries were sequenced on an Illumina HiSeq2500 (Illumina, San Diego, CA, USA) at the BGI Centre in Copenhagen, Denmark.

The study was approved by the Danish ethics committees and the Danish Data Protection Agency.

### 2. Data filtering

#### 2.1. Phenotype Assignment and Ancestry

Schizophrenia cases have been defined strictly as individual with diagnosis under criteria F20 under ICD-10, and controls as random sampled individuals without F20 diagnosis. European ancestry has been defined as having all 4 grandparents from Denmark, Scandinavia or at least Europe: the ancestry information has been used to define the parameters for PCA filtering (see below).

#### 2.2. Relatedness

The filtered variants dataset has been used in order to identify a subset of variants to be used for relatedness analysis. To this goal, SNPs only have been chosen with HWE p-value> 0.000001, MAF > 0.02, call rate >90% using PLINK 1.9. Kinship analysis has been performed using PLINK and individuals with a score >0.2 have been flagged as related. Based on this list of pairs of related individuals, a prioritisation on phenotype has been performed in order to select only one individual from the pair and exclude the other. Individuals with schizophrenia have been preferentially included, individuals with uncertain phenotype have been always excluded, when both individuals were cases with schizophrenia a random selection was made.

#### 2.3. Principal Component Analysis

Principal Component Analysis has been performed in HAIL, using a list of variants selected from the sample according to the following criteria: (a) variants have been filtered in HAIL with a call rate >= 0.9, bi-allelic only and with minor allele frequency >1%; (b) the variants have been then processed in PLINK for LD pruning (“indep-pairwise option”) using a window of 2000, step size of 20 variants and r2 threshold of 0.01.

Using the above selected list, the first 10 principal components have been calculated in HAIL and used to annotate the samples. The subsequent analysis has been performed in R. The European ancestry information (see above) has been used to select only the individuals within the 99% of the normal distribution of PC pairs of the individuals with EU ancestry. Individuals falling out of these defined limits were excluded from the analysis. Around 14% of the initial sample has been excluded. A further PCA analysis of the retained samples has been the performed in HAIL and we verified that no remaining variance in the PCs could be attributed to the ancestry: the new loadings have then been used as covariants for subsequent analyses.

#### 2.4. Variants and samples filtering

The analysis ready dataset has been prepared by excluding any variants with the following characteristics:

1. Any call with a depth a) less than 10 or b) greater than 1000;
2. Homozygous reference calls with a) GQ less than 25 or b) less than 90% reads supporting the reference allele;
3. Homozygous variant calls with a) PL(HomRef) less than 25 or b) less than 90% reads supporting the alternate allele;
4. Heterozygote calls with a) PL(HomRef) less than 25, b) less than 25% reads supporting the alternate allele, c) less than 90% informative reads (e.g. number of reads supporting the reference allele plus number of reads supporting the alternate allele less than 90% of the read depth), d) a probability of drawing the allele balance from a binomial distribution centered on 0.5 of less than 1e-9, or e) a location where the sample should be hemizygous (e.g. calls on the X chromosome outside the pseudoautosomal region in a male);
5. Any call on the Y chromosome outside the pseudoautosomal region on a sample from a female.

Following the application of these genotype filters, three call rate filters were used: first the removal of variants with a call rate below 90%, then the removal of samples with a call rate below 95%.

Samples with total URV counts exceeding the 95 percentile of the distribution in each dataset have been removed from the analysis-ready dataset. Supplementary Figure S1 shows the composition of the datasets at the different stages of filtering.

#### 2.5. Variant Annotation

Variants have been annotated using Variant Effect Predictor, and the Loftee plugin.

Ultra rare variants have been annotated according to the criteria described in Supplementary Figure 2, using Hail. Based on the above annotation, counts table have been produced using Hail.

### 3. Exome-wide analyses of ultra-rare variants

#### 3.1 Logistic regression analysis

We followed a similar approach as in Genovese et al. 2016, and therefore we have calculated a logistic regression on the schizophrenia status for the whole genome with the following model

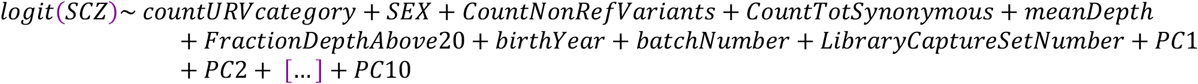

The Odds ratio and 95% confidence intervals for the enrichment in different URV categories of missense-constrained genes as shown in Figure S3.

#### 3.2 Meta-analysis of the three datasets

Once we obtained the significance estimates according to the approach described in 2.1, we used the R package “metagen” (https://cran.r-project.org/web/packages/metagen/index.html) in order to carry out the meta-analysis. We then plotted the fixed-effect results in each plot, as shown in Figure S3.

### 4. Gene-sets analyses of ultra-rare variants

Also in this step, we followed a similar approach as in Genovese et al. 2016, and therefore we have calculated a logistic regression on the schizophrenia status for the whole exome with the following model

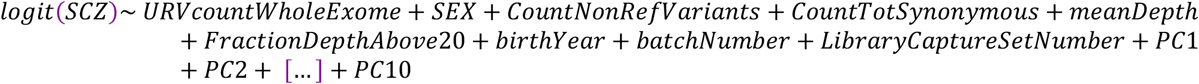

Then we have calculated the logistic regression for each specific gene-set on the genome-wide count of variants as follows

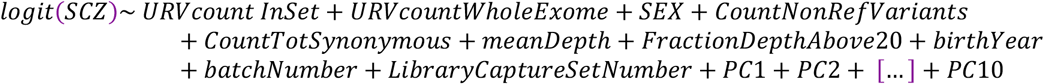

And in order to evaluate the effective contribution of the specific URV counts in the tested set, we have compared the 2 model with ANOVA and used its chi-square test p-value as adjusted p-value for our analysis.

Figure S4 shows the gene sets most significantly enriched genome-wide by dURVs, and comparison with enrichment of Synonymous URVs in the same sets.

### 5. Gene-level SKAT for rare variants

We used RAREMETAL (see: Feng, S., Liu, D., Zhan, X., Wing, M. K., & Abecasis, G. R. (2014). RAREMETAL: fast and powerful meta-analysis for rare variants. Bioinformatics, 30(19), 2828-2829. http://doi.org/10.1093/bioinformatics/btu367) in order to: 1. calculate summary statistics on the 2 single studies, and generate covariance matrices In this step the first 4 Principal Components (see above final PCA in each dataset) have been used as covariates; 2. combine the two studies and perform meta-analysis of single variants association and gene-collapsed methods (Burden, SKAT, VT, MB).

Only variants with the following annotations have been included in the counts: transcript_ablation, splice_donor_variant, splice_acceptor_variant, stop_gained, frameshift_variant, stop_lost, start_lost, initiator_codon_variant, missense_variant, protein_altering_variant, splice_region_variant, incomplete_terminal_codon_variant, mature_miRNA_variant, TFBS_ablation, TF_binding_site_variant.

Figure S5 shows a QQ-plot of the SKAT analysis performed with rare variants (i.e. MAF <0.05).

### 6. Gene-level SKAT for ultra-rare variants

Using RAREMETAL and the same approach as described in paragraph 4, we have performed the same analysis on ultra-rare variants.

Figure S6 shows the resulting QQ-plot of the SKAT performed with ultra-rare variants (see above, for criteria of annotation).

Supplementary Table 2 contains the full results from the RAREMETAl run.

### 7. extTADA Analysis

#### 7.1 Pre-analysis: identifying best clusters

As described by extTADA developers in Nguyen et al. 2017, we sought to identify subsets of our data where covariates had little effect on the results on the logistic regression of the URV counts.

Using the package mclust (https://cran.r-project.org/web/packages/mclust/index.html), we applied a parameterized Gaussian mixture model fitted by EM algorithm initialized by model-based hierarchical clustering using the matrix of covariates in the three different datasets.

We summarized the resuls using Bayesian Information Criterion (BIC) for the specified mixture models numbers of clusters, and we identified the most optimal clusters in each datasets.

Using this information, we partitioned the datasets and run on each partition the same logistic regression as described in 2.1 above, with and without covariates, to verify the correlation of the 2 analysis results.

Figure S7 shows regression analysis and resulting r2 values between p-values of logistic regression perform with covariates, and pvalues of logistic regression performed without covariates, within each of the clusters identified by multivariate cluster analysis of the datasets.

Given the above shown results, we used the identified partitions to carry out the following extTADA analysis.

#### 7.2 extTADA analysis

The extTADA analysis has been carried out as described in Nguyen et al. 2017, and for the de-novo counts and mutation rates we used the sample-adjusted ratios of mutation counts between 1,077 cases and 731 control presented in Nguyen work.

Figure S8 shows the pairs plot of the extTADA analysis, showing the estimates for pi0 and hypergamma on each of the dataset clusters identified in the clustering analysis.

### 8. Replication and combined p-values

### 8.1 Replication studies in UK samples

In order to replicate our results, the top 100 genes resulting from the extTADA analysis have been verified against the UK10K schizophrenia exome sequencing dataset (1,352 cases and 4,769 controls) and an Ion Torrent targeted sequencing study (5,207 cases and 4,991 controls) performed by Michael Owen, Michael O’Donovan, Elliot Rees and colleagues at Cardiff University. The dataset has been analysed using Firth’s penalized logistic regression model.

### 8.2 Combined analysis of results

Method for combining with formula form Jakob on effective counts The effective counts have been calculated as

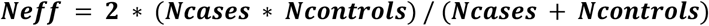

Where *Ncases* and *Ncontrols* are the total number of cases and controls dataset used for the combined analysis. This is the number of cases in a hypothetical sample if it is assumed to be balanced with respect to controls and having the same statistical power as the current study with *Ncases* and *Ncontrols*.

The above calculated effective counts have been used as weights, in order to combine the p-values of the different studies using the weighted sum z method implemented in the package *metap* (https://cran.r-project.org/web/packages/metap/index.html) and according to suggestions in Zaykin DV (2011). “Optimally weighted Z–test is a powerful method for combining probabilities in meta–analysis.” Journal of Evolutionary Biology, 24, 1836– 1841.

### 9. Gene-set enrichment of top extTADA results

We performed a gene-set enrichment of the extTADA results, for those genes with a posterior probability higher than 0.5.

In order to carry out the analysis, we have used the gene-sets described in Genovese et al. 2016, as well as the genes identified in Saeterstrom et al. 2019 (the full set of Autism Sequencing Consortium (ASC) genes, ASD-predominant, and genes for ASD with NDD).

The analysis has been performed with a hypergeometric test, using the function *phyper* from the stats package in R; p-values have been adjusted using the Benjamini & Hochberg method (1995).

Supplementary Table 3 reports the full results of this analysis, with FDR-corrected p-values.

## Footnotes

1 The iPSYCH2012 case-cohort sample: new directions for unravelling genetic and environmental architectures of severe mental disorders. (2018). The iPSYCH2012 case-cohort sample: new directions for unravelling genetic and environmental architectures of severe mental disorders., *23*(1), 6–14. http://doi.org/10.1038/mp.2017.196

2 Mors, O., Perto, G. P., & Mortensen, P. B. (2011). The Danish Psychiatric Central Research Register:. *Scandinavian Journal of Public Health*. http://doi.org/10.1177/1403494810395825

