## Supplementary Figures for "Meta-analysis of Scandinavian Schizophrenia Exomes"

### Supplementary Figures - Meta-analysis of Scandinavian Schizophrenia Exomes

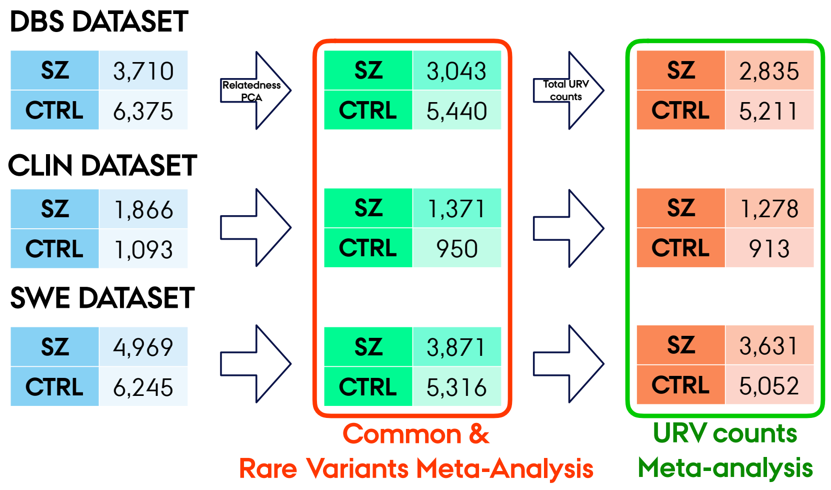


Figure S1: Dataset composition and filtering steps

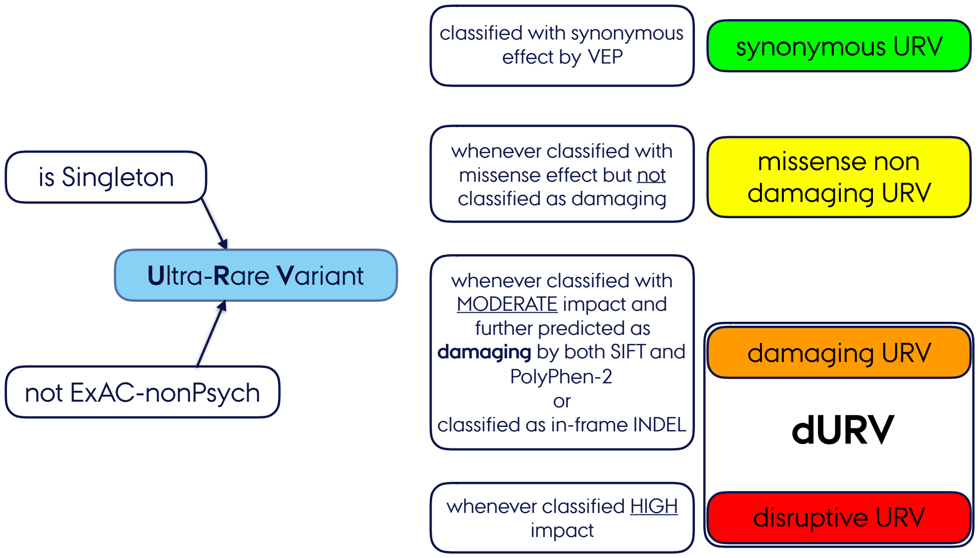


Figure S2: criteria for annotation of ultra-rare variants

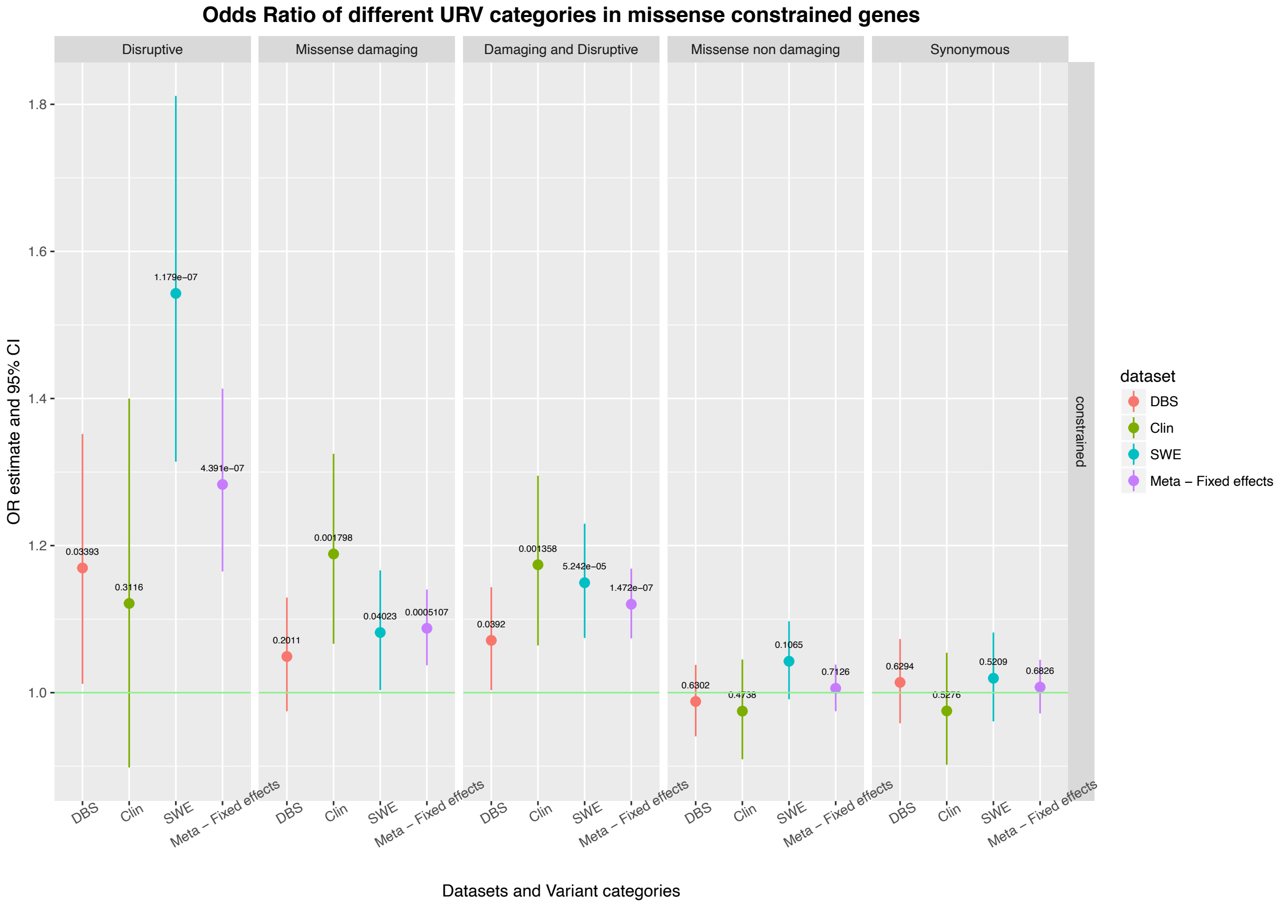


Figure S3: Odds ratio and 95% confidence intervals for the enrichment in different URV categories of missense-constrained genes.

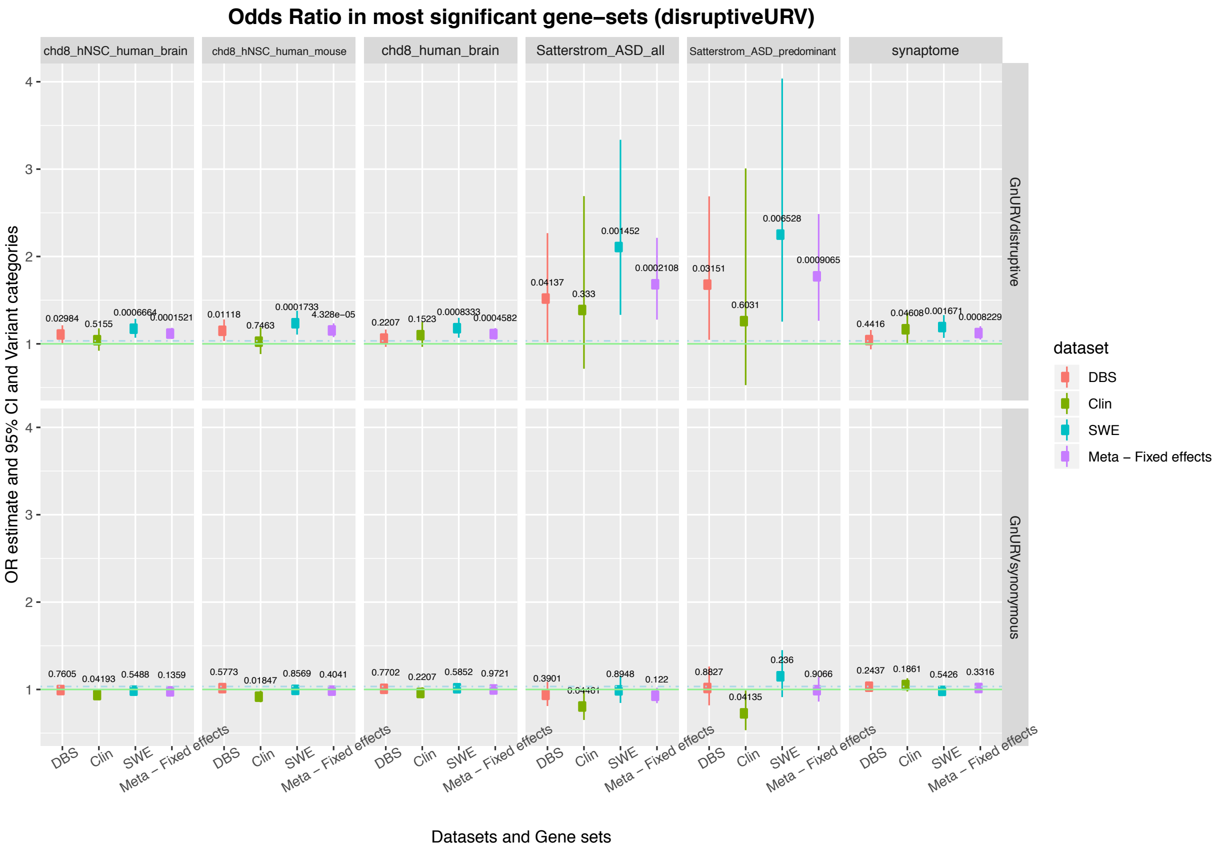

Figure S4: Gene sets most significantly enriched genome-wide by dURVs, and comparison with enrichment of Synonymous URVs in the same sets.

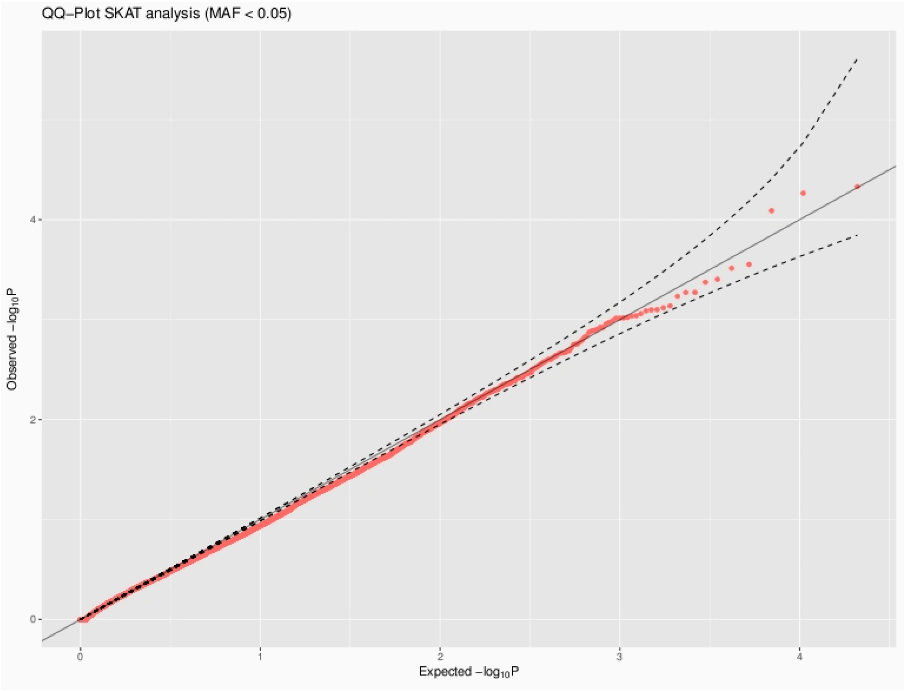


Figure S5: QQ-plot of the SKAT analysis performed with rare variants (i.e. MAF <0.05).

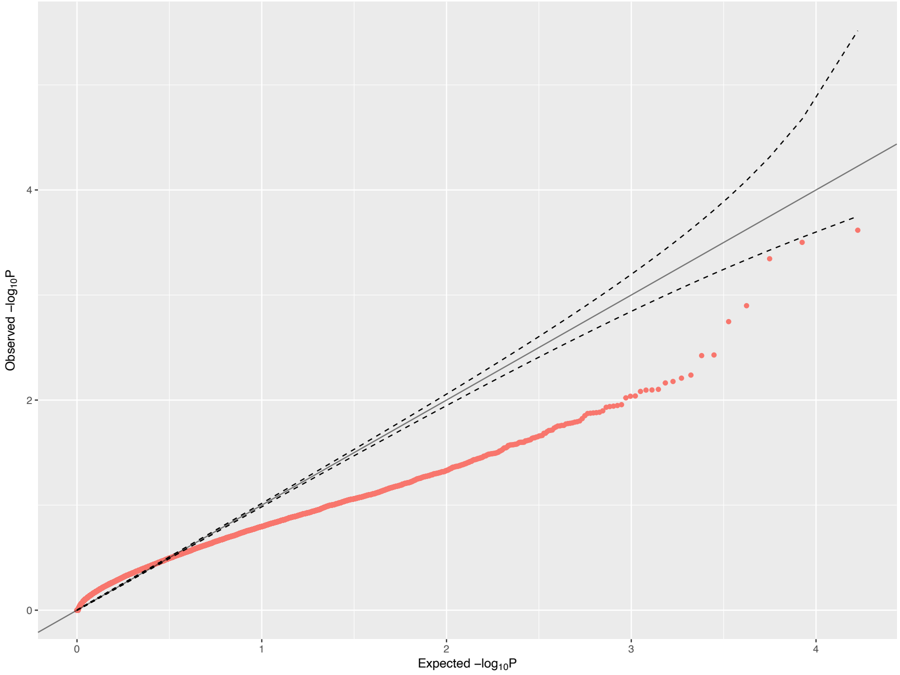


Figure S6: QQ-plot of the SKAT performed with ultra-rare variants (see above, for criteria of annotation)

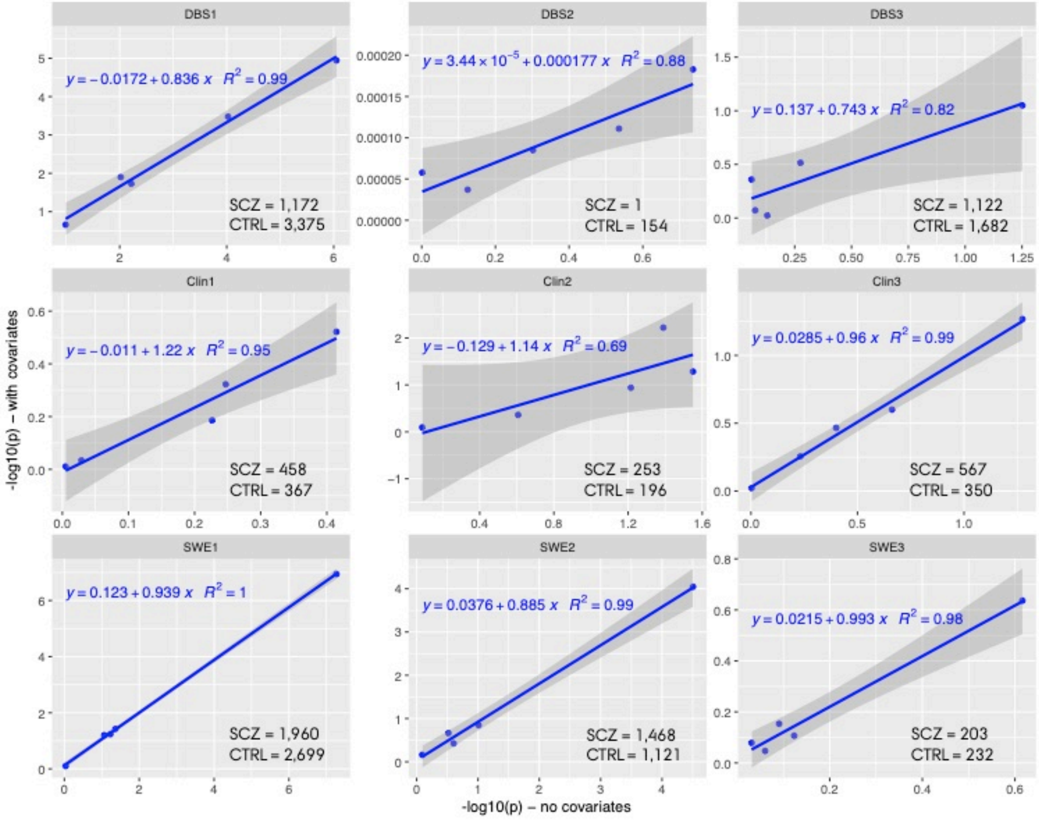


Figure S7: regression analysis and resulting r2 values between pvalues of logistic regression perform with covariates, and pvalues of logistic regression performed without covariates, within each of the clusters identified by multivariate cluster analysis of the datasets.

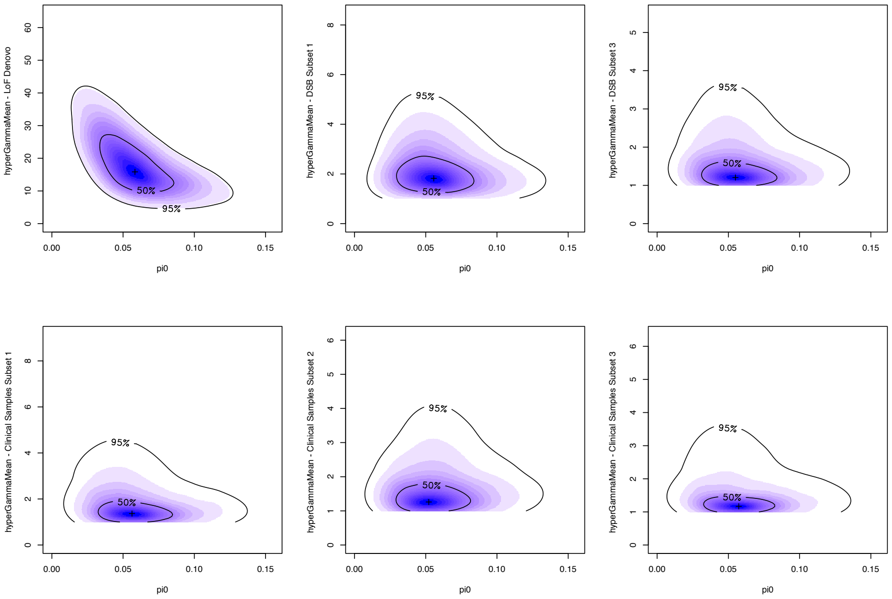

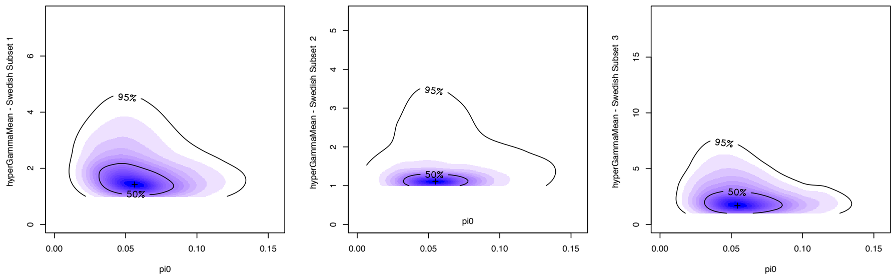


Figure S8: Pairs plot of the extTADA analysis, showing the estimates for pi0 and hypergamma on each of the dataset clusters identified in the clustering analysis.
